# Resting heart rate rapid reduction by moderate exercise evolutionarily encoded

**DOI:** 10.1101/155663

**Authors:** Gábor Pavlik, Eva Bakács, Eszter Csajági, Tibor Bakács, Judit Noe, Robert Kirschner

## Abstract

**Background:** Global physical inactivity pandemic is responsible for more than 5 million deaths annually through its effects on non-communicable diseases. This requires urgent intervention.

**Objective:** To investigate associations of physical activity with cardiovascular fitness in a cross-sectional retrospective observational study. Data were collected for 21 years from 2530 healthy volunteers and athletes representing the entire spectrum of physical activity from the totally inactive sedentary persons to the highly trained national athletes.

**Methods:** Simple echocardiographic parameters of cardiovascular fitness were analyzed. Cardiac fitness was characterized by reduced resting heart rate, increased relative left ventricular muscular mass, improved left ventricular diastolic function and peak exercise oxygen consumption.

**Results:** We found that even moderate exercise is associated with improved cardiac fitness. The largest improvement of fitness was observed between the inactive and the least active group, whereas fitness decreased in the highly trained national athletes enduring up to 20 training hours per week.

**Conclusions:** Our finding that moderate exercise is associated with positive changes in sedentary persons makes sense only in the light of evolution. Human endurance running performance capabilities that emerged ~2 million years ago are evolutionary coded and seems to be awakened even by moderate exercise. This finding would help physicians to encourage patients for doable and sustainable behavioral change who are currently inactive and find physical exercise intimidating. (Word count: 218)

**Abbreviations:**

CV: (cardiovascular)
CVD: (cardiovascular disease)
CVH: (cardiovascular health)
HD: (heart disease)
BSA: (body surface area)
LV: (left ventricular)
RHR: (Resting Heart Rate)

**Key Points:** This cross-sectional retrospective observational echocardiographic study of 2530 healthy volunteers and athletes representing the entire spectrum of physical activity from the totally inactive sedentary persons to the highly trained national athletes found that it is possible to experience cardiovascular benefits soon after the sedentary persons become physically active. This makes sense only in the light of evolution. With increasing performance level cardiovascular fitness is increased up to a point but then decreased in highly trained national athletes.

The non-invasive and simple echocardiographic test could be used to monitor exercise induced positive changes. This would help physicians in their efforts to promote the expansive benefits of exercise in all spectrums of society and encourage patients for doable and sustainable behavioral change.

## 1 Introduction

Physical inactivity became a global pandemic responsible for over 5 million deaths annually through its effects on multiple non-communicable diseases require urgent multidisciplinary health response (1). Not unexpectedly, exercise capacity is a more powerful predictor of cardiovascular (CV) disease and mortality than any other established CV risk factors (2) (3).

Aerobic exercise induces beneficial physiological left ventricular (LV) remodeling, numerous progressive cardiac adaptations, which are collectively termed the “athlete’s heart” (4). Despite nonlinear anatomic and electrical remodeling, the athlete’s heart retains normal or supernormal myocyte function (5). Structurally, all four heart chambers increase in volume with increased wall thickness, resulting in greater cardiac mass due to increased myocardial cell size (6). The cardiovascular adaptation for generating a large and sustained increase in cardiac output during prolonged exercise includes a 10-20% increase in cardiac dimensions (7). Changes of echocardiographic parameters (circumferential shortening velocity, rel. cardiac output at rest) have also been indicated as characteristic of the athlete’s heart (8, 9). Consistent with this, exercise training has become a class I recommendation in all major international guidelines as an evidence-based treatment of chronic heart failure (10).

While more than 100 papers demonstrated that moderate exercise is capable to improve cardiorespiratory fitness (CRF), the minimum intensity of physical activity that is associated with favorable body composition and CRF remains unknown. A new study found that in mid-childhood a higher intensity of physical activity was necessary to confer benefits to CRF than to improve body composition, but both associations were ultimately characterized by a dose-dependent phenomenon (11). The authors, therefore, concluded that displacing any lower-for-higher intensity activity in childhood may be an important first-order public health strategy.

In the above context our current study is encouraging because we demonstrate in more than 2500 healthy adult volunteers and athletes that regardless of how inactive you may be, it is possible to experience cardiovascular benefits soon after becoming physically active. By measuring the Resting Heart Rate (RHR) and simple echocardiographic parameters we show that the largest improvement of fitness following exercise was observed between the sedentary persons and the least active leisure group. In a PubMed search (as of April 2^nd^, 2017) we found no other paper in which the entire spectrum of physical activity from the totally inactive sedentary adults to the highly trained national athletes enduring up to ~20 gruelling training hours per week have been compared by echocardiography. Therefore, our findings are not only robust but novel as well, which has public health significance as it would help physicians to persuade sedentary patients to get active and engage in health-enhancing moderate physical activity. Such a way, they could encourage their patients to use their body “in the way it was designed” (12).

## 2 Methods

### 2.1 Healthy Volunteers and Athletes

Data of 2530 healthy volunteers and athletes (1617 men, 913 women, between 18 – 42 years of age) were analyzed in this cross-sectional retrospective observational study ^b^. Examinations were conducted during 1994-2015 at the Department of Health Sciences and Sports Medicine of the University of Physical Education, which is a National Institute recruiting athletes, but also physically active and inactive healthy volunteers from the entire country. Therefore, the participants collected for 21 years can be considered representative for the Hungarian population. Our primary goal was to collect sufficient amount of data for an analysis to determine the basic echocardiographic parameters that measure the effect of physical activity on the heart as we described earlier (13). This study meets the ethical standards of the IJPM (14).

The following six categories of performance levels were considered: inactive persons (Non-A); leisure time athletes (Leisure-A); lower class athletes (Lower cl-A); second class athletes (2nd cl-A); first class athletes (1st cl-A) and athletes of national teams (National-A) (see Table 1). The following athletes participated in the study: endurance athletes (343 men vs. 121 women), dry-land and water ball-game-players (423 vs. 274), sprinters-jumpers (61 vs. 45), power athletes (147 vs. 36), artistic sports (17 vs. 60) and sports students (18 vs. 25). Data in the figure and table are combined groups. VO_2_ max was determined only in a subset of the subjects (N=1192). Number of participants with missing data for each variable of interest is listed in Table 2. A questionnaire was completed first documenting the training habits/ history, medical/family history, whereas blood pressure, body height and weight were recorded.

**Table 1.**
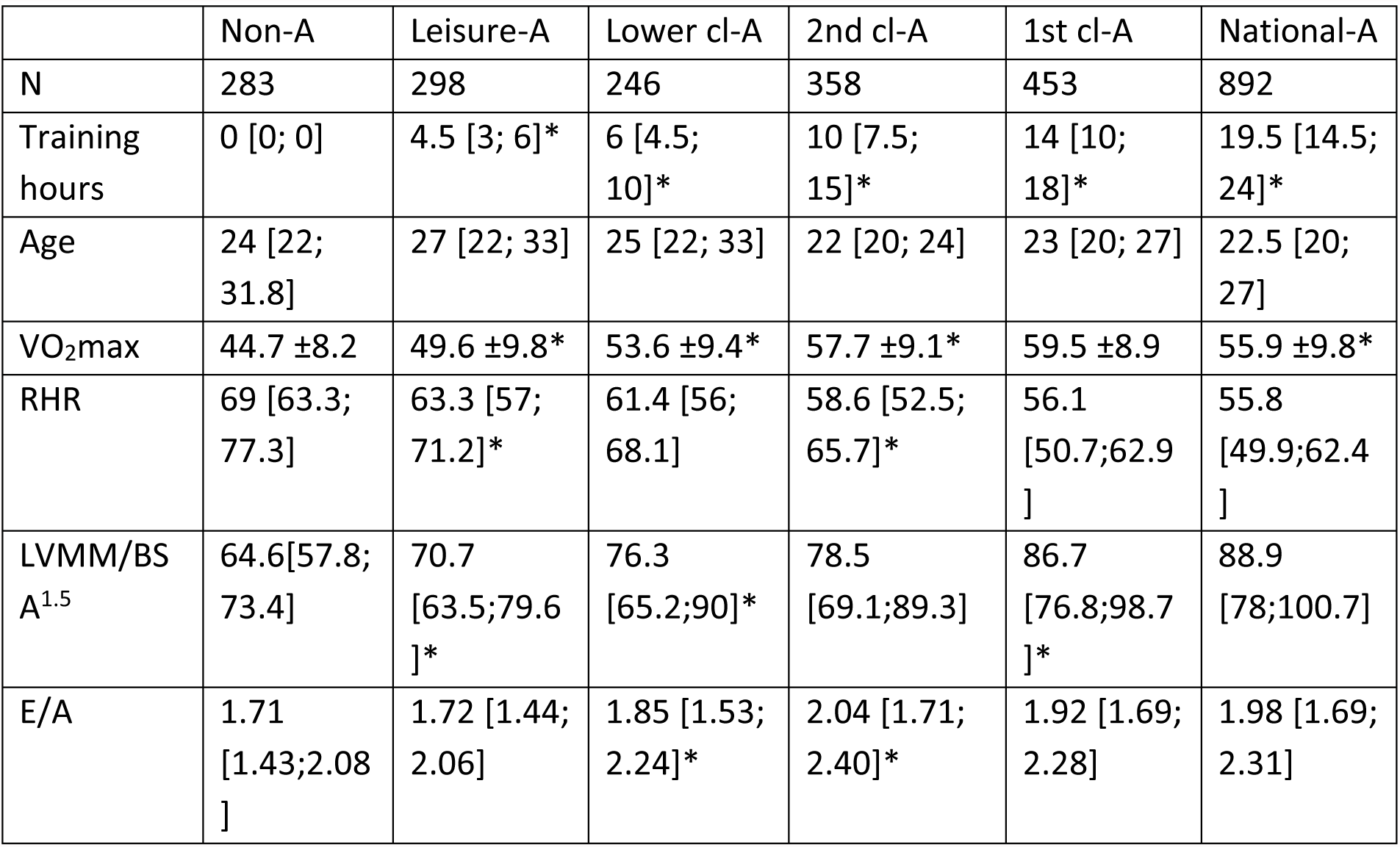
Characteristics of the study

**Table 2.** Number of participants with missing data for each variable of interest

|  | Non-A | Leisure-A | Lower cl-A | 2nd cl-A | 1st cl-A | National-A |
| --- | --- | --- | --- | --- | --- | --- |
| N | 283 | 298 | 246 | 358 | 453 | 892 |
| VO<sub>2</sub>max | 168 | 184 | 254 | 165 | 220 | 347 |
| RHR | 1 | 1 | 0 | 0 | 2 | 2 |
| LVMM/BSA<sup>1.5</sup> | 1 | 0 | 0 | 0 | 1 | 1 |
| E/A | 6 | 4 | 31 | 4 | 16 | 98 |

### 2.2 Echocardiography

2D-guided M-mode and Doppler-echocardiography were carried out by two devices. In the first period (1994-2009) measurements were made by a Dornier AI 4800 echocardiograph 2.5 MHz transducer, during the second period by a Philips HD15 echocardiograph (1–5 MHz, Koninkiljke Philips, NV, USA) in left lateral decubitus position. From the parasternal view, 2D-guided M mode recordings, left ventricular (LV) end diastolic septal (IVSd), posterior wall thickness (PWTd), and end diastolic dimension (LVIDd) were acquired. All examinations were carried out by the same physician (G.P.). Each parameter was estimated on 3–5 heart cycles to minimize intra-observer variability. The averages were used in further analysis. Relative left ventricular muscular mass (Rel.LVMM) was calculated by the cubing method, values were referred to the indexed body surface area (BSA), so that the numerator and the denominator have the same power (8, 15). Diastolic function was characterized by the quotient of the early (E) and late (A) peak diastolic velocities (E/A). Resting Heart Rate (RHR) was measured on the Doppler echocardiographic curves.

### 2.3 Statistics

Statistical analysis was carried out using the 5.3 Version of SigmaStat for Windows (SPSS Inc., Chicago, USA). The normality of the distribution of different statistical variables was analyzed with the Kolmogorov-Smirnoff test. Variables are reported as mean ± SD if they passed the test of normality (only VO_2_ max variable). Otherwise, they are reported as median with 25^th^ and 75^th^ percentiles shown in brackets [quartile]. While VO_2_ max variable passed either the Kolmogorov-Smirnoff or the equal of variances test, one-way analysis of variance was used for the analysis of the difference of mean of VO_2_ max between different performance levels. An overall significance (P<0.001) was established for rejecting the null hypotheses for VO_2_ max that the six treatment groups were not different. Accordingly, pair wise differences between the groups were revealed by using Tukey’s method of multiple comparisons. For all other variables, the Kruskal-Wallis analysis of variance for nonparametric variables was used for calculating the difference of median in the different study groups. An overall significance (P<0.001) was established for rejecting the null hypotheses for all these variables that the six treatment groups were not different. Pair wise differences between the groups were revealed by using Dunn’s method of multiple comparisons. P≤0.01 was considered significant. Missing data were handled by SigmaStat for Windows.

## 3 Results

Performance levels of the athletes are correlated with their weekly training hours that differed significantly from the neighboring group in each performance class (Table 1). With increasing performance level, VO_2_ max elevated significantly (P<0.01) until a plateau was reached in the 2nd cl-A and the 1st cl-A class groups (N.S.). Then, VO_2_ max decreased significantly (P<0.01) between the 1st cl-A and National-A classes forming an inverse U-shaped curve (Figure 1A). A steep, significant (P<0.01) reduction of Resting Heart Rate (RHR) was apparent, when the Non-A and Leisure-A groups were compared (Figure 1B). Despite of the fact that the athletic performance level increased substantially, the RHR decreased further only moderately. In fact, there was no significant difference between the RHR of Leisure-A and Lower cl-A groups (N.S.). Furthermore, there was no significant difference between the National-A and the 1st cl-A groups (N.S.) or between the 2nd cl-A and 1st cl-A athletes (N.S.). Rel.LVMM increased with increasing performance level (Figure 1C). However, the difference was much more marked between the Non-A and Leisure-A groups (P<0.01), and between the Leisure-A and Lower cl-A groups (P<0.01), than among the different higher class athletic groups (N.S., except between the 2nd cl-A and the 1st cl-A groups (P<0.01)). E/A quotient was higher in athletes (Lower cl-A, 2nd cl-A, 1st cl-A and National-A groups), than in the Non-A and Leisure-A groups (Figure 1D). With increasing performance level an inverse U-shaped curve was observed, the peak of which was in the 2nd cl-A group. The differences between the Leisure-A and Lower cl-A groups and between the Lower cl-A and 2nd cl-A groups were significant (P<0.01). The highest E/A quotient was found in the 2nd cl-A group, which decreased again in the 1st cl-A group (N.S.) and then rose again in National-A group (P<0.01).

**Figure 1 A, B, C, D.**
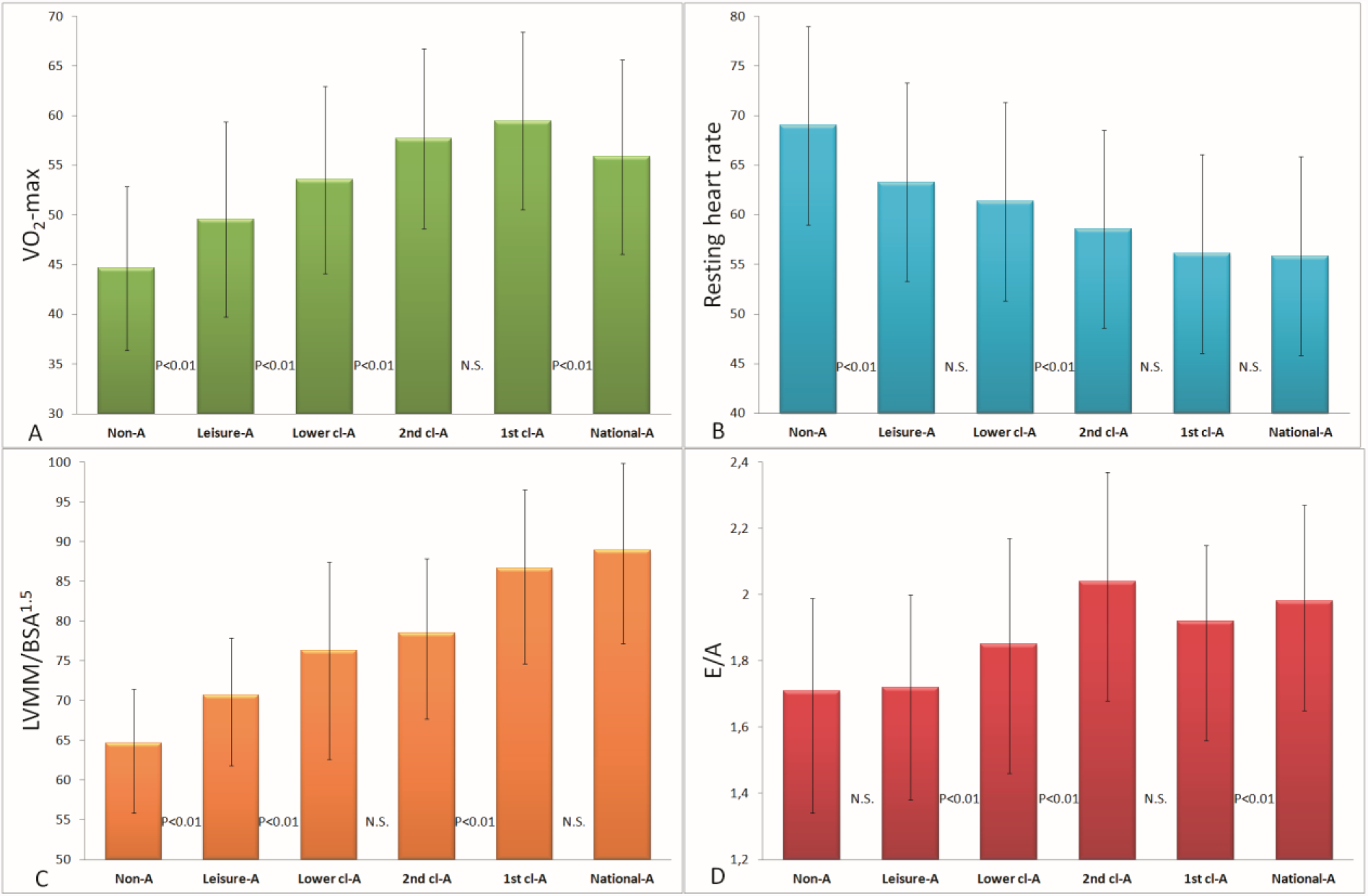
Peak exercise oxygen consumption (VO_2_ max) (A), resting heart rate (RHR) (B), relative left ventricular hypertrophy (LVMM) (C), left ventricular diastolic function (E/A) (D).

## 4 Discussion

Our results indicate that even moderate exercise is associated with positive changes in the characteristics of the athlete’s heart, which improves cardiac fitness and possibly lowers the risk of CV diseases. Three characteristics of the heart were analyzed in this study, each representing a different physiologic property. Rel.LVMM is a morphologic characteristic, which markedly differed between the 1st cl-A and 2nd cl-A groups and between the 2nd cl-A, Lower cl-A and Non-A groups, whereas it reached a peak in the 1st cl-A and National-A groups. The consensus view is that following physical training LV cells are undergoing a physiological adaptive hypertrophy and will have better pumping function resulting in higher maximum cardiac output during peak level exercise. It should be emphasized that the LV hypertrophy is considered a risk factor in the general medical praxis, since it is the manifestation of several cardiovascular abnormalities (cardiomyopathy, hypertension, etc.). However, several reports described the morphologic and functional differences between physiologic and pathologic LV hypertrophy (8, 16-19).

### 4.1 Physiologic and Pathologic LV Hypertrophy

One important difference between the physiologic and pathologic LV hypertrophy reported in our previous studies is that the former can only be induced by exercise at young age. Above 40-45 years, the LV hypertrophy is a dangerous risk factor, and cannot be considered a physiologic characteristic of the physically active person (13). Another basic difference between the physiologic and pathologic LV hypertrophy is that the latter is always associated with an increased stiffness of the heart manifested by an impaired diastolic function while it is not an issue in the athlete’s heart. In our current study the diastolic function was characterized by the E/A quotient, the ratio of the peak velocities of the early to late phase of the diastolic filling. The E/A quotient was not impaired in our athletic persons, and the highest value was shown in the 2nd cl-A groups. The better relaxation ability may increase the pumping function by a higher contribution of the Frank-Starling mechanism. It should also be noted that very intensive training (performed in the 1st cl-A and National-A groups, respectively), which induces additional LV hypertrophy, may decrease the CVH benefit obtained with less exercise.

The resting bradycardia of the athletes is induced mostly by elevated parasympathetic activity. It is beneficial because the diastolic period is significantly expanded during bradycardia. Considering that the coronary circulation of the LV is penetrable only during diastole, a lower RHR has marked benefits. We should therefore emphasize that following moderate exercise the largest decrease of RHR was observed between the sedentary persons (Non-A) and the least active leisure group (Leisure-A) in our study.

### 4.2 Evolutionary Perspective

Our finding that even moderate exercise is associated with improved CVH makes sense only in the light of evolution. Human endurance running performance capabilities probably emerged about 2 million years ago in order to help meat-eating hominids to compete with other carnivores (20). Although humans no longer need to run for food, the capacity and proclivity to run marathons (42.2 km) is the modern manifestation of this trait. Since the average finish time in marathons hardly changes by age (e.g. see New York City Marathon: Average Finish Times by Age Group^c^), it was claimed that one cannot find any other field of athletic endeavor where sixty four year olds compete with nineteen year olds (20). According to Dr. Bramble, “We’re a machine built to run – and the machine never wears out…” consequently, “…humans really are obligatorily required to do aerobic exercise in order to stay healthy,…”.^d^

### 4.3 Doing Little Is Better Than Doing Nothing

However, in order to meet even a simplified aerobic exercise challenge, the physicians should wage a determined campaign, similar to that of the anti-smoking campaigners. Since the 1960s they have been compelled to challenge the perception that the smokers’ behavior is commonplace and integral to everyday life by revealing that the cigarette is dangerous, addictive, and deadly. Today, similarly to the anti-smoking campaigners, the physicians should explain their patients that they are capable to reactivate a genetically encoded running ability with relatively little physical effort. This is consistent with the study of Lee et. al, who demonstrated that running even 5 to 10 min/day at slow speeds is associated with markedly reduced risks of death from all causes and cardiovascular disease (21). The good news therefore is that the mortality benefits of light jogging may motivate healthy but sedentary individuals to begin and continue running for health benefits as a practical, achievable, and sustainable goal. For successful aging even a little physical activity is good. Doing little is better than doing nothing (22). This argument is also supported by a recent systematic meta-analysis that included 174 articles. (23) Continuous risk curves for the associations between total physical activity and risk of breast cancer, colon cancer, diabetes, ischemic heart disease, and ischemic stroke events showed that major gains occurred at lower levels of activity (up to 3000-4000 metabolic equivalent (MET) minutes/week), while the same amount of increase yielded much smaller returns at higher levels of activity.

### 4.4 Is It Possible to Have Too Much of a Good Thing?

Notwithstanding, it is still an unanswered question whether it is possible to have too much of a good thing. The Copenhagen City Heart Study in 1,098 healthy joggers and 3,950 healthy non-joggers suggested, for example, a U-shaped association between all-cause mortality and dose of jogging as calibrated by pace, quantity, and frequency of jogging (24). This is consistent with our observations. Specifically, the VO_2_ max values and the E/A ratios formed inverse U-shaped curves with increasing performance level where the highest values were observed in the 1st cl-A and 2nd cl-A groups, respectively (see Figures 1A and 1D). This is “good news” to encourage people because moderate exercise is a doable and sustainable objective. All the more so, since regardless of how inactive you may be, it is possible to experience cardiovascular benefits soon after becoming physically active (25).

In order to encourage achievable and sustainable behavioral change, Blair et al. (26) reviewed the physical activity recommendations as to how much and what intensity physical activity should be done. They emphasized that the consensus public health recommendation of 30 min of moderate intensity physical activity/d on ≥5 d/wk was largely directed at the 40–50 million US adults who are sedentary and who account for much of the public health burden of chronic disease. Blair et al., argue that even if the lower dose of exercise may be insufficient to prevent unhealthful weight gain for some, it should be recommended because data suggest that relatively small changes in activity or fitness on the part of sedentary persons might produce large reductions in disease risk at the population level. Blair’s concept is confirmed by many studies. Screening 835 reports in the PubMed and Embase databases, totaling 122417 participants, with a mean follow-up of 9.8+/-2.7 years and 18122 reported deaths (14.8%), Hupin at al (27) found that even 15 minutes a day is technically enough in order to reduce one’s risk of all-cause mortality. Importantly, a U shaped association was observed between the exercise capacity and arrhythmia in more than a million men at a median age of 18.2 years during a median follow-up of 26.3 years (28).

### 4.5 To Overcome the Study Limitations a Testable Hypothesis Is Proposed

A main limitation that our study is a retrospective analysis. Therefore, the trends observed should be confirmed by a prospective observational cohort study examining the effects of moderate physical exercise on heart rate reduction and on left ventricular remodeling by echocardiography. We predict that selective lowering of resting heart rate with moderate physical exercise will improve cardiovascular outcomes. The risk reduction capability of physical exercise can be estimated by a comparison with the results of the BEAUTIFUL and SHIFT randomized placebo-controlled studies, which assessed the effect of heart-rate reduction by the selective sinus-node inhibitor ivabradine on outcomes in heart failure (29, 30). These studies demonstrated that the risk of primary composite endpoint events increased by 3% with every beat increase from baseline heart rate and 16% for every 5-bpm increase; furthermore, that selective lowering of heart rates with ivabradine improved cardiovascular outcomes.

## 5 Conclusion

Physicians must redouble prevention efforts for the CVH by explaining their patients that the cardiac fitness can be improved by moderate physical activity. The patients’ efforts can be monitored by echocardiography, which does not require a sophisticated laboratory. Our evidence-based observation that moderate physical exercise is associated with significant improvements of cardiac fitness could help physicians to encourage patients for doable and sustainable behavioral change. Clinicians should therefore promote the expansive benefits of exercise in all spectrums of society, be it the casual exerciser, the sedentary individual or those with established CV disease (3). The non-invasive echocardiographic test monitoring exercise induced positive changes would help physicians in their efforts. (word count: 2997)

## Acknowledgment

Gábor Pavlik, Eszter Csajági and Robert Kirschner had full access to all of the data in the study and take responsibility for the integrity of the data and the accuracy of the data analysis. The authors gratefully thank Dr J. Mehrishi, PhD, Cantab, FRCPath of Cambridge, United Kingdom, for reviewing of manuscript and editing assistance. The authors also thank Dr. Duck-chul Lee, Assistant Professor, Department of Kinesiology, Iowa State University, USA, for his useful comments. Grant support: none.

## Conflicts of interest

none.

## Footnotes

a Quotes from Dr. Bramble, in Christopher McDougall: *Born to Run*; ps. 173; 240.

b This study was approved by the Semmelweis University, Regional and Institutional Committee of Science and Research Ethics; approval number: 121/2010.

c http://www.runtri.com/2010/11/new-york-city-marathon-average-finish.html

d Quotes from Dr. Bramble, in Christopher McDougall: *Born to Run*; ps. 173; 240.

## References

1. Sallis JF, Bull F, Guthold R, Heath GW, Inoue S, Kelly P, et al. Progress in physical activity over the Olympic quadrennium. Lancet. 2016.

2. Myers J, Prakash M, Froelicher V, Do D, Partington S, Atwood JE. Exercise capacity and mortality among men referred for exercise testing. N Engl J Med. 2002;346(11):793–801.

3. Otto CM. Heartbeat: highlights from this issue. Heart. 2015;101(10):739–41.

4. Fernandes T, Barauna VG, Negrao CE, Phillips MI, Oliveira EM. Aerobic exercise training promotes physiological cardiac remodeling involving a set of microRNAs. Am J Physiol Heart Circ Physiol. 2015;309(4):H543–H52.

5. Paterick TE, Gordon T, Spiegel D. Echocardiography: profiling of the athlete’s heart. J Am Soc Echocardiogr. 2014;27(9):940–8.

6. Wilson MG, Ellison GM, Cable NT. Basic science behind the cardiovascular benefits of exercise. Heart. 2015;101(10):758–65.

7. Sharma S, Merghani A, Mont L. Exercise and the heart: the good, the bad, and the ugly. Eur Heart J. 2015;36(23):1445–53.

8. Pavlik G, Major Z, Varga-Pinter B, Jeserich M, Kneffel Z. The athlete’s heart Part I (Review). Acta Physiol Hung. 2010;97(4):337–53.

9. Pavlik G, Major Z, Csajagi E, Jeserich M, Kneffel Z. The athlete’s heart Part II Influencing factors on the athlete’s heart: Types of sports and age (Review). Acta Physiol Hung. 2013;100(1):1–27.

10. Adams V, Niebauer J. Reversing heart failure-associated pathophysiology with exercise: what actually improves and by how much? Heart Fail Clin. 2015;11(1):17–28.

11. Collings PJ, Westgate K, Vaisto J, Wijndaele K, Atkin AJ, Haapala EA, et al. Cross-Sectional Associations of Objectively-Measured Physical Activity and Sedentary Time with Body Composition and Cardiorespiratory Fitness in Mid-Childhood: The PANIC Study. Sports Med. 2017;47(4):769–80.

12. Lee IM, Shiroma EJ, Lobelo F, Puska P, Blair SN, Katzmarzyk PT. Impact of Physical Inactivity on the World’s Major Non-Communicable Diseases. Lancet. 2012;380(9838):219–29.

13. Pavlik G, Olexo Z, Osvath P, Sido Z, Frenkl R. Echocardiographic characteristics of male athletes of different age. Br J Sports Med. 2001;35(2):95–9.

14. Harriss DJ, Atkinson G. Ethical Standards in Sport and Exercise Science Research: 2016 Update. Int J Sports Med. 2015;36(14):1121–4.

15. Pavlik G, Olexo Z, Frenkl R. Echocardiographic estimates related to various body size measures in athletes. Acta Physiol Hung. 1996;84(2):171–81.

16. Cheng TO. Hypertrophic cardiomyopathy vs athlete’s heart. Int J Cardiol. 2009;131(2):151–5.

17. Hildick-Smith DJ, Shapiro LM. Echocardiographic differentiation of pathological and physiological left ventricular hypertrophy. Heart. 2001;85(6):615–9.

18. Pelliccia A, Maron BJ. Outer limits of the athlete’s heart, the effect of gender, and relevance to the differential diagnosis with primary cardiac diseases. Cardiol Clin. 1997;15(3):381–96.

19. Richey PA, Brown SP. Pathological versus physiological left ventricular hypertrophy: a review. J Sports Sci. 1998;16(2):129–41.

20. Bramble DM, Lieberman DE. Endurance running and the evolution of Homo. Nature. 2004;432(7015):345–52.

21. Lee DC, Pate RR, Lavie CJ, Sui X, Church TS, Blair SN. Leisure-time running reduces all-cause and cardiovascular mortality risk. J Am Coll Cardiol. 2014;64(5):472–81.

22. Gebel K, Ding D, Bauman AE. Physical Activity and Successful Aging-Reply: Even a Little Is Good. JAMA Intern Med. 2015;175(11):1863–4.

23. Kyu HH, Bachman VF, Alexander LT, Mumford JE, Afshin A, Estep K, et al. Physical activity and risk of breast cancer, colon cancer, diabetes, ischemic heart disease, and ischemic stroke events: systematic review and dose-response meta-analysis for the Global Burden of Disease Study 2013. BMJ. 2016;354:i3857.

24. Schnohr P, O’Keefe JH, Marott JL, Lange P, Jensen GB. Dose of jogging and long-term mortality: the copenhagen city heart study. J Am Coll Cardiol. 2015;65(5):411–9.

25. Chomistek AK, Henschel B, Eliassen AH, Mukamal KJ, Rimm EB. Frequency, Type, and Volume of Leisure-Time Physical Activity and Risk of Coronary Heart Disease in Young Women. Circulation. 2016;134(4):290–9.

26. Blair SN, LaMonte MJ, Nichaman MZ. The evolution of physical activity recommendations: how much is enough? The American Journal of Clinical Nutrition. 2004;79(5):913S–20S.

27. Hupin D, Roche F, Gremeaux V, Chatard JC, Oriol M, Gaspoz JM, et al. Even a low-dose of moderate-to-vigorous physical activity reduces mortality by 22% in adults aged >/=60 years: a systematic review and meta-analysis. Br J Sports Med. 2015;49(19):1262–7.

28. Andersen K, Rasmussen F, Held C, Neovius M, Tynelius P, Sundstrom J. Exercise capacity and muscle strength and risk of vascular disease and arrhythmia in 1.1 million young Swedish men: cohort study. BMJ. 2015;351:h4543.

29. Bruguera CJ, Varela A. Role of heart rate in cardiovascular diseases: how the results of the BEAUTIFUL study change clinical practice. Am J Cardiovasc Drugs. 2009;9 Suppl 1:9–12.

30. Swedberg K, Komajda M, Bohm M, Borer JS, Ford I, Dubost-Brama A, et al. Ivabradine and outcomes in chronic heart failure (SHIFT): a randomised placebo-controlled study. Lancet. 2010;376(9744):875–85.

